# Conformational and interface variability of bivalent RNF4-SIM diSUMO3 interaction

**DOI:** 10.1101/2021.06.17.448718

**Authors:** Alex Kötter, Henning D. Mootz, Andreas Heuer

## Abstract

SUMO targeted ubiqutin ligases (STUbLs) like RNF4 or Arkadia/RNF111 recognize SUMO chains through multiple SUMO interacting motifs (SIMs). Typically, these are contained in disordered regions of these enzymes and also the individual SUMO domains of SUMO chains move relatively freely. It is assumed that binding the SIM region significantly restricts the conformational freedom of SUMO chains. Here, we present the results of extensive molecular dynamics simulations on the complex formed by the SIM2-SIM3 region of RNF4 and diSUMO3. Though our simulations highlight the importance of typical SIM-SUMO interfaces also in the multivalent situation, we observe that frequently other regions of the peptide than the canonical SIMs establish this interface. This variability regarding the individual interfaces leads to a conformationally highly flexible complex. Even though this is in contrast to previous models of the RNF4 SUMO chain interaction, we demonstrate that our simulations are clearly consistent with previous experimental results.

## Introduction

The post-translational modification by the small ubiquitin-like modifier SUMO (SUMOylation) is implicated in a wide range of cellular processes.^1–3^ On the molecular level, it is often the ability of SUMO to non-covalently bind coregulatory proteins, that is critical for these processes.^4,5^ Typically, this non-covalent interaction is mediated by a SUMO interacting motif (SIM) of the coregulatory protein.

The highly similar paralogs SUMO2 and SUMO3 are efficiently assembled into polymeric chains via SUMOylation of Lys11 upon triggering the SUMO response. As SUMO chains provide an array of interaction sites for SIMs, the conjugation of SUMO chains is a distinct signal for proteins with multiple SIMs through multivalency.^6^ This mechanism can, for example, be observed in the arsenic-based treatment of acute promyelocytic leukemia (PML). Here, the PML protein is polySUMOylated upon treatment with arsenic trioxide. Recognition of the SUMO chains by the E3 ubiquitin ligase RNF4 in turn leads to ubiquitination and subsequent degradation of the PML protein.^7,8^

RNF4 is a member of the family of SUMO-targeted ubiquitin ligases (STUbLs) and, besides Arkadia/RNF111, the only known human STUbL.^9,10^ STUbLs contain multiple SIMs to connect the SUMOylation and ubiquitination pathways. RNF4, for example, contains four SIMs that are part of its disordered N-terminal region.^11–14^ The involvement of STUbLs in processes like the DNA damage response and chromatin organization is a highly active research field. ^6,15^

Multiple experimentally determined structures established the typical picture of the SIM-SUMO interface.^16–20^ Upon binding SUMO, the short hydrophobic SIM forms a *β*-sheet that aligns with the second *β*-sheet of SUMO and the hydrophobic side chains of the SIM occupy the hydrophobic SIM binding groove of SUMO. It seems obvious to assume that such interfaces are also present in multivalent complexes formed by SIM-regions of STUbLs and SUMO chains. Here, the investigation of structural properties is, however, complicated by the fact that these SIM-regions are typically disordered, the SUMO chains do not adopt a defined tertiary structure and the individual SIM-SUMO interaction is typically relatively weak.

Despite the flexibility of the constituents, it is assumed that complexes formed by SIM-regions of STUbLs and SUMO chains adopt a defined structure. For example, Kung et al. modeled the structure of the tetravalent complex formed by the RNF4-SIM region and tetraSUMO2 based on experimental results of the corresponding monovalent complexes^11^ and Xu et al. used NMR to solve the bivalent structure of the RNF4-SIM2-SIM3 diSUMO3 complex.^13^ In fact, this is the only experimentally determined structure of a SUMO chain with SIM binding partners. The structure appears questionable regarding the details of the individual SIM-SUMO interfaces, as it does not show the typical binding mode observed for monovalent SIM-SUMO complexes. ^21^

Another somewhat puzzling result was reported by Keusekotten et al., who found the affinity between tetraSUMO2 and mutants of the RNF4-SIM region with only two intact SIMs remaining, to be the same as for the wild type RNF4-SIM region, although it would be expected that more SIM-SUMO interactions lead to higher affinity binding. ^12^

Given the poor understanding of SUMO chain recognition by polySIM proteins and the conformational states before and after binding, it is highly desirable to obtain a clearer picture of the interface between SUMO chains and the SIM region of RNF4, also because next to its own physiological importance, the RNF4-SUMO interaction serves as a reference in the investigation of similar systems. ^22–25^

Here, we present large-scale replica exchange molecular dynamics simulations of the RNF4-SIM2-SIM3 diSUMO3 complex. Our simulations reveal the variable role of SIM-SUMO interfaces. These are surprisingly found both in binding modes highly similar to the known experimental structures of monovalent complexes and in those that are atypical with residues outside the classical SIM sequences establishing the interaction with the SIM binding groove. We furthermore observe that both the unbound SUMO3 dimer as well as the bound bivalent complex do not adopt a defined structure, but rather represent an ensemble of conformations in which the positions of the SUMO monomers relative to one another appear highly flexible, a behavior that is reminiscent of complexes involving diUbiquitin. ^26^

Due to the microscopic information, available in simulations, our conclusions go beyond the previous interpretation of experimental results. Our results from computer simulations turn out to be compatible with key experimental results on this system, however. As shown in this work, this encompasses, e.g., the influence of binding RNF4-SIM2-SIM3 on the conformation of diSUMO3 observed in FRET experiments ^27^ and on the relative motion of the SUMO3 units observed in NMR experiments by Xu et al. ^13^

## Results

### The SIM2, SIM3-diSUMO3 complex does not adopt a defined structure

Xu et al. reported the structure of the complex formed by the SIM2-SIM3 region of RNF4 and diSUMO3 (PDB: 2mp2).^13^ As input for our simulations, we used the best representative conformer of PDB file 2mp2. In contrast to Xu et al., however, we used a linear instead of an isopeptide linked SUMO3 dimer, which is an establied and commonly used surrogate in the investigation of multivalent SIM-SUMO interaction. ^7,12,22^ We used replica exchange MD simulations (REST2^28^) to efficiently sample the configuration space of the SIM2-SIM3-bound diSUMO3 (30 replicas, 3 *μ*s simulation time per replica) and the unbound diSUMO3 (17 replicas, 1 *μ*s simulation time per replica). For the analysis, we saved configurations every 1 ns.

In figure 1 A we show 100 randomly picked structures sampled during the simulations of the bound and free systems. To visualize the relative position of the individual SUMO3 domains, we aligned the structures with respect to the backbone of the distal and proximal units of the SUMO3 dimer (i.e. N-terminal and C-terminal SUMO3, N-SUMO3 and C-SUMO3).

**Figure 1:**
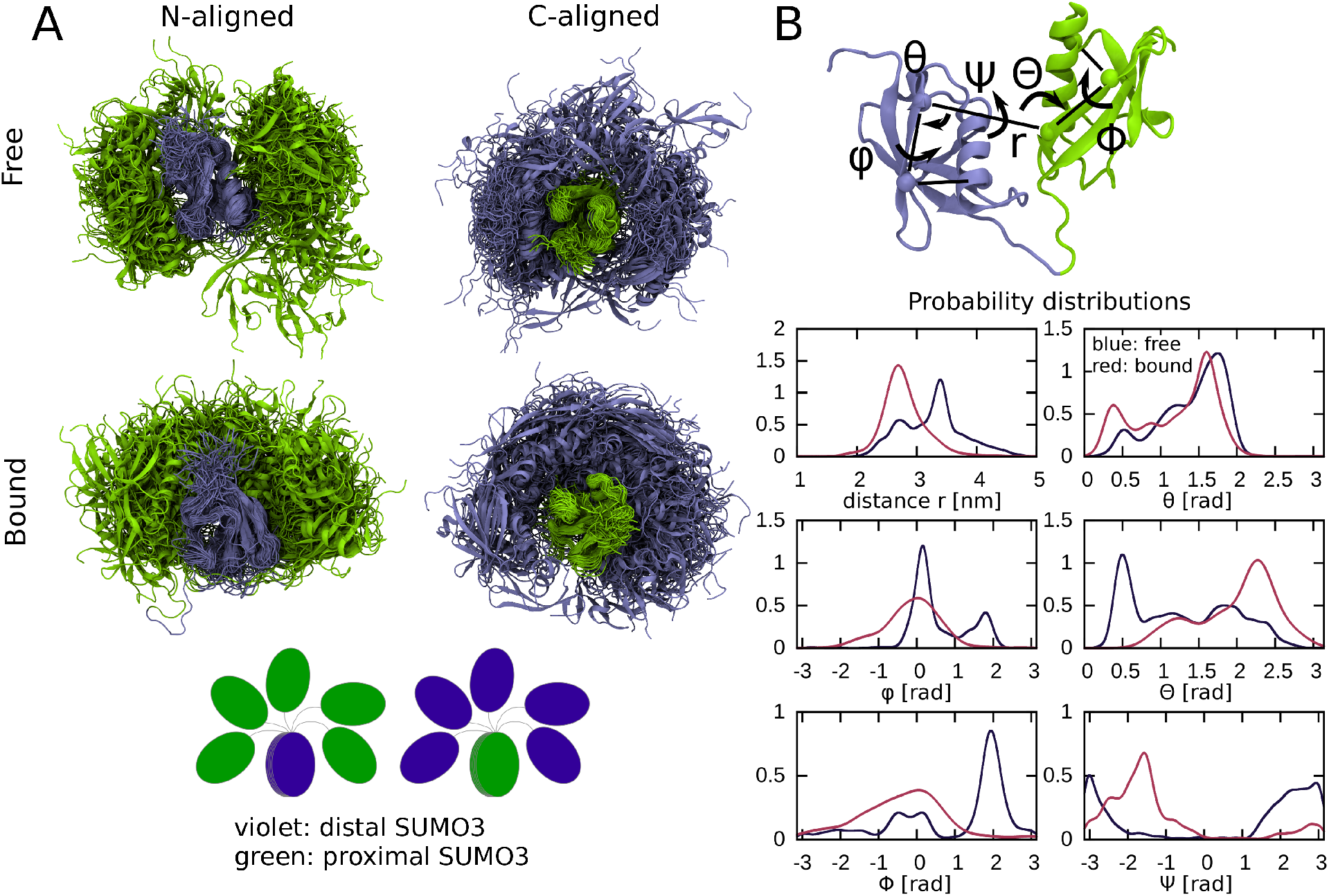
A: 100 randomly chosen structures sampled during our REST2 simulations of the free (top) and bound system (bottom). Distal SUMO3 (N-SUMO3) and proximal SUMO3 (C-SUMO3) are depicted in violet and green, respectively. The left and right panels show structures that are aligned with respect to distal and proximal SUMO3, respectively. B: Definition and probability distributions of collective variables that are used to monitor the diSUMO3 conformation. The variables are defined in terms of residues Ile29 and Ile33 at the N-terminal and C-terminal ends of the second *β*-sheet and Cys47 at the C-terminal end of the *α*-helix of SUMO3.

In agreement with previous experiments, ^11–13^ the free SUMO3 dimer does not adopt a defined tertiary structure in our simulations. Also in the bound state, no defined structure is adopted and the conformational freedom of the SUMO3 dimer is not strongly restricted.

Nevertheless, we observe that binding the SIM2-SIM3 peptide alters the populations of possible diSUMO3 conformations. Consider, for example, the position of the proximal SUMO3 with respect to the SIM binding groove of the distal SUMO3. In the free state, the SIM binding groove is clearly visible in figure 1 A. In the bound state it is, however, hidden behind multiple SUMO3 structures.

The influence of binding on the diSUMO3 conformation can also be captured by a set of collective variables. Their definition, and distributions estimated from the simulations of the diSIM-bound and free SUMO3 dimer can be found in figure 1 B. We observe broad distributions for all variables both in the free and in the bound state. Some of the distributions are also multimodal, indicating a complex conformational behavior. Furthermore, we clearly observe that binding of the SIM2-SIM3 peptide alters the distributions of the variables and therefore the conformational behavior of the SUMO3 dimer.

A conformational change of a SUMO2 dimer induced by binding the SIM2-SIM3 peptide has been observed previously in FRET experiments. ^27^ Here, fluorophores were attached to positions Arg61 of each SUMO2 unit. Upon binding the SIM2-SIM3 peptide, the distance between the fluorophores increased. Figure S1 shows the distribution of the Arg60-Arg60 distance in bound and free state, respectively, as estimated from our simulation data (Arg60 of SUMO3 corresponds to Arg61 of SUMO2). In agreement with the experiments, binding the SIM2, SIM3 peptide leads to a larger Arg60-Arg60 distance in our simulations, thus further corroborating the reliability of the simulation.

Based on similar rotational autocorrelation times of the SUMO3 units in the bound state, Xu et al. concluded that these move as an essentially rigid entity in solution, in contrast to the situation in the free state. Based on this observation, they further concluded that the conformational freedom of the SUMO3 dimer is severely limited when bound to the SIM2-SIM3 peptide.^13^ To investigate the relative motion of the SUMO3 monomers in the bound and free state, we conducted further ordinary MD simulations because dynamical information is lost through the Monte Carlo moves in REST2 simulations. We initiated a single additional simulation for the free system and two additional simulations for the bound system, starting in different initial structures.

In figure S2, we show time series of the collective variables defined in figure 1 B in the these simulations. We observe that the relative motion of the SUMO3 monomers is severely slowed down by binding the SIM2-SIM3 peptide. In the bound state, the time series exhibit long plateaus with minor fluctuations. In contrast, in the free state, we observe strong fluctuations also on short time scales for multiple collective variables. This observation clearly agrees with the NMR measurements of Xu et al. which show the relative motion of the SUMO3 monomers is restrained by binding the SIM2-SIM3 peptide. Apparently we cannot conclude from this, however, that there is a single favored conformation of the SUMO3 dimer.

### Individually, the SIM2 and SIM3 regions bind in multiple interfaces

To analyze the sampled structures with respect to the individual SIM-SUMO interfaces, we considered the two sets of structures that were aligned either with respect to the distal or proximal SUMO3 and clustered these sets with respect to the atom positions of the SIM2 and SIM3 regions, respectively. This way, we sorted the structures into different groups based on the individual interface between a single SUMO3 domain and SIM2 and SIM3, respectively. The sequence of the SIM2-SIM3 peptide can be found in figure 2 A.

**Figure 2:**
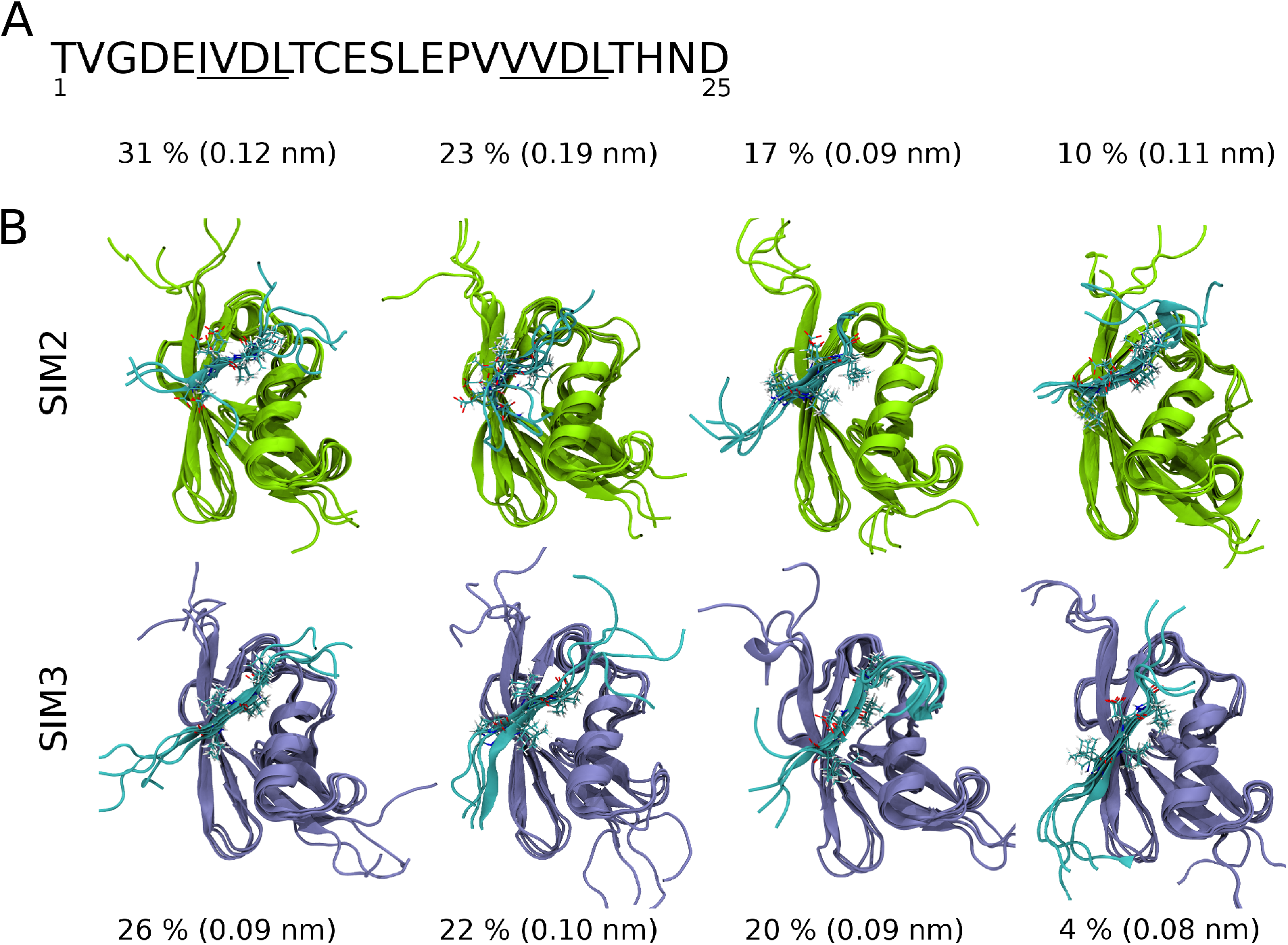
A: Sequence of the SIM2-SIM3 peptide, SIM2 and SIM3 are underlined. B: Representative structures for the clusters of the SIM2 and SIM3 regions. In all cases the most representative structure and three median structures are shown. The frequencies of the clusters are given as percentages and the RMSD of the median structure to the most representative structure is given in parenthesis. The structures were aligned with respect to the SUMO3 that is bound to the respective SIM region (proximal SUMO3 for SIM2, top, and distal SUMO3 for SIM3, bottom).

In total, 86 % of the sampled structures were assigned to a SIM2 cluster and 88 % of the sampled structures were assigned to a SIM3 cluster. The remaining structures are considered noise. Representative structures for the most frequently visited clusters with a probability of at least 10 % can be found in figure 2. For the SIM3 region, we include one additional cluster with a probability of only 4 % because this cluster consists of structures that exhibit the classical parallel SIM3-SUMO interface.

We defined the most representative structure of a cluster as that structure that minimizes the root mean squared deviation (RMSD) to all other structures of the cluster. In addition to this most representative structure, we also give three median structures in figure 2 (see also the methods section). Table 1 summarizes the results of the cluster analysis with a focus on the residues that occupy the SIM binding groove and the orientation of the SIM region with respect to the second *β*-sheet of SUMO3. Note that the residues which most frequently occupied the SIM binding grooves in our simulations are the same that were most strongly affected by binding SUMO2/3 chains in the NMR experiments by Xu et al.^13^ and Kung et al.^11^

**Table 1:** Results of the cluster analysis: The column “structure” gives the amino acids that occupy the SIM binding groove and the orientation of the SIM region with respect to the SUMO3 domain it interacts with. Furthermore, the table shows the probability of the clusters, estimated from all sampled structures (*p_tot_*) and from the first, second and third 1000 sampled structures (*p*_1*st*_, *p*_2*nd*_, *P*_3*rd*_). The clusters are sorted according to the observed probabilities.

| cluster | structure | fig | $p_{tot}$ | $p_{1st}$ | $p_{2nd}$ | $p_{3rd}$ |
| --- | --- | --- | --- | --- | --- | --- |
| SIM2 1 | Aps4, Ile6, para | 2 | 0.31 | 0.42 | 0.33 | 0.18 |
| SIM2 2 | Val7, para | 2 | 0.23 | 0.16 | 0.27 | 0.25 |
| SIM2 3 | Val7 Leu9, para | 2 | 0.17 | 0.14 | 0.27 | 0.11 |
| SIM2 4 | Val7 Leu9, anti | 2 | 0.10 | 0.04 | 0.06 | 0.20 |
| SIM2 5 | helix, para | S3 | 0.05 | 0 | 0.04 | 0.10 |
| SIM3 1 | Val17 Val19, anti | 2 | 0.26 | 0.05 | 0.21 | 0.52 |
| SIM3 2 | Val17 Val19, para | 2 | 0.22 | 0.08 | 0.37 | 0.22 |
| SIM3 3 | Val19 Leu21, anti | 2 | 0.20 | 0.35 | 0.13 | 0.11 |
| SIM3 4 | Leu21, anti | S3 | 0.06 | 0.17 | 0.02 | 0 |
| SIM3 5 | Val18 Asp20, anti | S3 | 0.05 | 0.13 | 0.03 | 0 |
| SIM3 6 | Pro16 Val18, anti | S3 | 0.05 | 0.01 | 0.09 | 0.05 |
| SIM3 7 | Val19 Leu21, para | 2 | 0.04 | 0.07 | 0.02 | 0.03 |

The four clusters for each SIM that are shown in figure 2 account for 81 % of the sampled structures in case of SIM2, and for 72 % of the sampled structures in case of SIM3. For the SIM2 region, we observed one additional cluster with a probability of 5 % and for the SIM3 region, we observed three additional clusters with probabilities of 5 to 6 %. Representative structures for these clusters can be found in figure S3.

Except for the second cluster of the SIM2 region, the most important clusters comprise of structures that are highly similar and exhibit SIM-SUMO interfaces, in which the SIM region forms a *β*-sheet and two residues of the SIM occupy the SIM binding groove of SUMO3 (figure 2).

While usually two hydrophobic residues occupy the SIM binding groove, in cluster 1 of the SIM2 region, it is not the two expected residue Ile6 and Leu8, but rather Asp4 and Ile6 that occupy the groove. A similarly unusual SIM-SUMO interface has been reported for the complex formed by Thymine-DNA Glycosylase and SUMO3, where valine and glutamate occupy the SIM binding groove (PDB: 1wyw).^29^ However, in the latter case the SIM-SUMO interaction occurs through an intramolecular arrangement and therefore might be more prone to engage unusual SIM sequences.

Cluster 2 of the SIM2 region comprises of structures that are less similar among each other than in case of the other clusters. Essentially, the only common structural element in this cluster is that Val5 occupies the position close to the first helix turn in the SIM binding groove.

Clusters 1 and 2 of the SIM3 region comprise of structures in which residues PVVV act as SIM. Recently, a similar SIM-SUMO interface was observed in the complex formed by the viral STUbL ICP0 and SUMO2. Here, the SIM comprises of residues PIVI, with Ile2 and Ile4 occupying the SIM binding groove.^30^ Clusters 3 and 4 of the SIM3 region correspond to the expected SIM3-SUMO interfaces in antiparallel and parallel orientation, respectively.

In table 1 we give the probability estimates from three blocks of the simulation. Furthermore, we give the simulation time dependence of the probability estimates in figure S4. The clusters shown in figure 2 are visited in all parts of the simulation. Therefore all these clusters are relevant in the sense that their observation was not a non-equilibrium effect. The simulation time dependence of the probability estimates and the fluctuations between the different parts of the simulation indicate, however, that the probability estimates did not converge and that consequently they cannot be used as quantitative estimates of the experimental probabilities of the clusters.

In the most representative conformer in the ensemble of structures reported by Xu et al. (PDB: 2mp2),^13^ the system adopts an overall antiparallel interaction pattern, with SIM2 interacting with the proximal SUMO3 and SIM3 interacting with the distal SUMO3. During our simulation, this interaction pattern does not change. Xu et al, however, also reported structures with an overall parallel arrangement, in which SIM2 interacts with the distal SUMO3 and SIM3 interacts with the proximal SUMO3, suggesting that flipping the orientation of the SIM2-SIM3 peptide is possible.

We therefore performed additional REST2 simulations in the alternative overall parallel arrangement and analyzed the impact of this parameter on the individual SIM-SUMO interactions in the same way as for the previous simulation. In the overall parallel configuration, less structures could be assigned to clusters of the individual SIM-SUMO interfaces (72 % and 77 % of all sampled structures for SIM2 and SIM3, respectively). Representative structures of these clusters and a summary of the cluster analysis are given in figure S9 and table S3, respectively. The simulation time dependence of their probability estimates is given in figure S10. Beside the individual SIM-SUMO interfaces, the overall orientation of the complex also influences the conformation of the SUMO3 dimer (see figure S11)

In the overall parallel arrangement, typical SIM-SUMO interfaces have lower probability. Furthermore, in 9 % of all sampled structures in the parallel arrangement, the SIM2 region did not interact with SUMO3. In most of these structures, SIM3 forms a typical parallel SIM3-SUMO interface (see also figure S12). Apparently this interface is able to stabilize the monovalent complex in the ensemble of bivalent structures. These observations indicate that the overall antiparallel arrangement of the most representative conformer of the NMR ensemble is more relevant than the overall parallel arrangement.

In the following, we will therefore focus on the analysis of the overall antiparallel arrangement of the bivalent diSIM-diSUMO complex. Additional information for the parallel arrangement can be found in the supporting information.

### Correlations among the SIM-SUMO interfaces

To analyze correlations between the SIM2 and SIM3 interfaces, we compared the actually observed probability for the interface combination *i/j*, i.e. the situation where the SIM2 region falls in cluster *i* and the SIM3 region falls in cluster *j*, with the product of the individual probabilities of the clusters *i* and *j* (table 2). That means, we compare P(SIM2 binds in cluster *i* and SIM3 binds in cluster *j*) with P(SIM2 binds in cluster *i*) × P(SIM3 binds in cluster *j*). Table 2 shows this analysis only for individual clusters of the SIM2 and SIM3 regions that we observed with a probability of at least 10 %. Similar data for the remaining clusters with lower probabilities can be found in tables S1 and S2. Additionally, we also provide the same analysis for the simulation of the overall parallel configuration in table S4.

**Table 2:** Probabilities of interface combinations. The product of the probabilities of the individual clusters is given in parenthesis. We underlined some interface combinations with relatively high probability. In parenthesis we indicate the orientation of the interface (parallel (p) or antiparallel (a)).

|  | SIM3 noise | SIM3 1 (a) | SIM3 2 (p) | SIM3 3 (a) |
| --- | --- | --- | --- | --- |
| SIM2 noise | 0.04 (0.02) | 0.01 (0.04) | 0.01 (0.03) | 0.05 (0.03) |
| SIM2 1 (p) | 0.03 (0.04) | 0.005 (0.08) | <u>0.08</u> (0.07) | <u>0.12</u> (0.06) |
| SIM2 2 (p) | 0.01 (0.03) | <u>0.12</u> (0.06) | 0.01 (0.05) | 0.02 (0.04) |
| SIM2 3 (p) | 0.02 (0.02) | 0 (0.05) | <u>0.12</u> (0.04) | 0.003 (0.03) |
| SIM2 4 (a) | 0.01 (0.01) | <u>0.07</u> (0.03) | 0 (0.02) | 0.01 (0.02) |

Our analysis shows that most interface combinations are visited only rarely or not at all, leaving only a handful of high probability interface combinations that account for 50 % of the sampled structures. In the simulation of the overall parallel configuration, we observe a similar behavior (table S4).

The time dependence of the probability estimates of the interface combinations can be found in figure S4. We observe that the high probability combinations 1/3, 2/1, 3/2 and 1/2 are visited early in the simulation. The combination of two typical antiparallel SIM-SUMO interfaces (4/1) is visited only after two thirds of the simulation. This indicates that its probability may be underestimated.

Structures with interface combination 3/2 and with combination 4/1 correspond to conformations with two typical SIM-SUMO interfaces in parallel and antiparallel orientation, respectively. Additional conformational or steric demands arising in the bivalent complex would be one possible explanation for the unusual bonding modes of SIM2 and SIM3 in the NMR structure of the bivalent complex (PDB: 2mp2^13^). Representative structures with interface combinations 3/2 and 4/1, that can be found in figure S5 indicate, however, that the system is able to bind in two typical SIM-SUMO interfaces without any obvious steric or conformational problems for either diSUMO3 or the SIM2-SIM3 peptide. From the knowledge of monovalent SIM-SUMO complexes, one might have expected these to be the dominating interface combinations in addition to other possible combinations of typical SIM-SUMO interfaces that we, however, only observed with very little probability if at all. Through visual inspection of these structures with low probability interface combinations (e.g. combination 3/3) no obvious reasons for their low probability could be identified (see figure S6 for representative structures).

### Interface combination and diSUMO3 conformation are highly correlated

In the following, we systematically analyze the correlation of the interface combination and the diSUMO3 conformation. We focus on the four interface combinations 3/2, 1/3, 2/1 and 4/1 because they are visited with relatively high probability and contrast structures with two typical SIM-SUMO interfaces and structures with a single typical SIM-SUMO interface.

We used the set of structures that was aligned with respect to the backbone of the distal SUMO3. To capture the relative position of the proximal SUMO3 in a set of structures that exhibits a certain interface combination, we determined the most representative structure of the proximal SUMO3 in this set. This most representative structure minimizes the backbone RMSD to all structures of the proximal SUMO3 in the set (for more information, see the methods section). In figure 3 A, we show the most representative and nine median structures for the four investigated interface combinations. Figure 3 B gives the average RMSD to to the most representative structure of the proximal SUMO3 for the interface combinations and, for reference, also for all bound structures.

**Figure 3:**
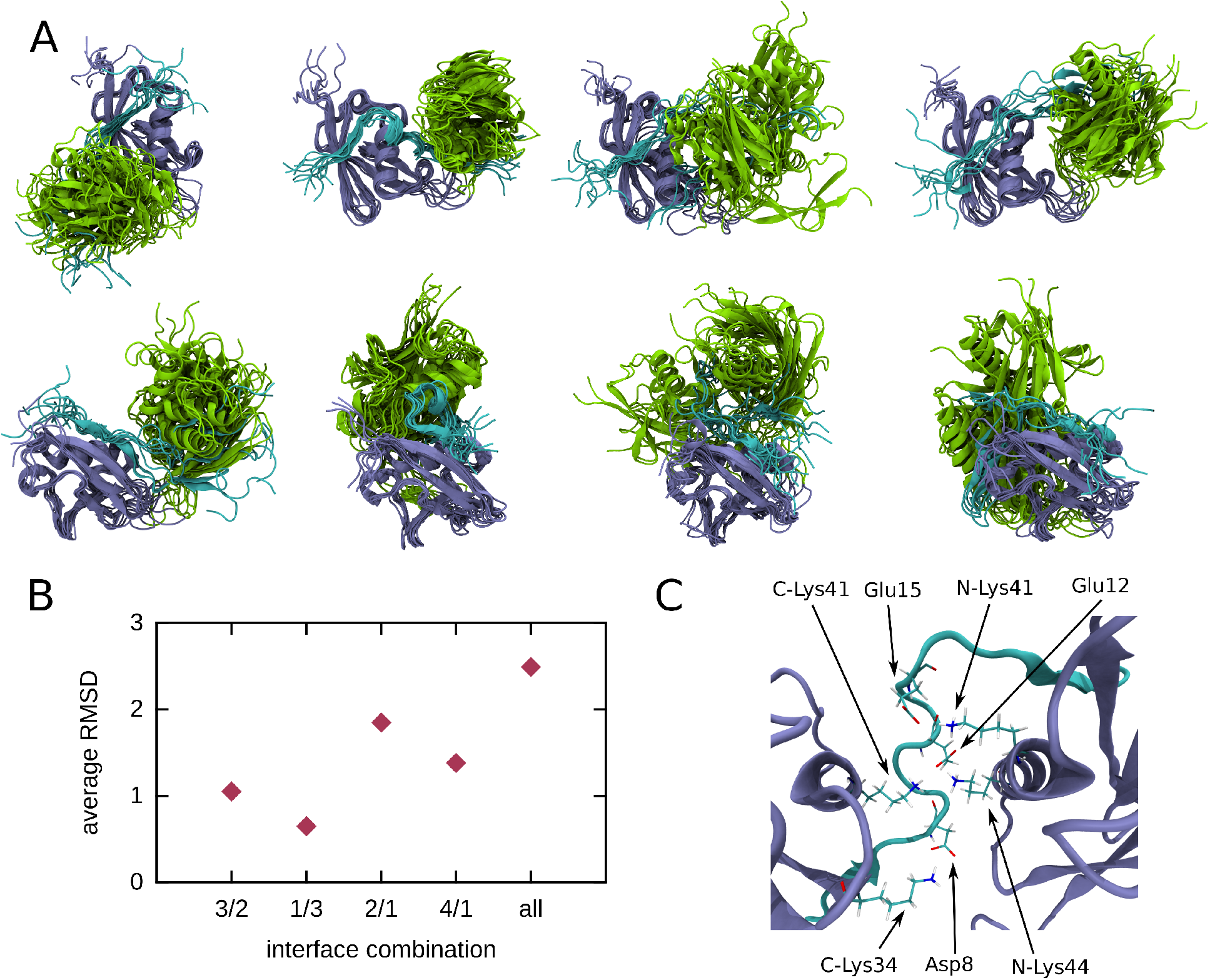
Figure A shows representative conformations of the complex when bound in interface combinations 3/2, 1/3, 2/1, and 4/1, respectively. We aligned the system with respect to the backbone of the distal SUMO3, top and bottom rows show the same structures from different perspectives. Distal SUMO3, proximal SUMO3 and SIM2-SIM3 peptide are depicted in violet, green and cyan, respectively. Figure B shows the average RMSD of the backbone of the proximal SUMO3 to the most representative structure for the interface combinations and for all bound structures. Figure C shows a structure of the complex with interface combination 1/3 and the electrostatic interaction between residues of the linker between the SIMs and SUMO3 residues highlighted.

When considering only structures that exhibit the same interface combination, the relative conformation of the SUMO3 units is strongly restrained. This is clearly in contrast to the situation in figure 1 A, where the relative conformation of the SUMO3 units in randomly chosen structures was considered. The diSUMO3 conformation is most restricted in interface combination 1/3, followed by combinations 3/2 and 4/1 and least restricted for combination 2/1. Furthermore, the conformation of diSUMO3 differs between the interface conformation. Most clearly, this can be observed for the relative position of the SUMO3 units in interface combination 3/2 and 1/3, respectively. For example, for interface combinations 1/3 and 4/1, the relative position of the SUMO3 units is more similar, their relative orientation, however, differs greatly, as indicated by orientation of the *α*-helix of the proximal SUMO3.

This behavior can also be captured in terms of the collective variables defined in figure 1 B. In figure S7, we give the distributions of these collective variables estimated from structures with the considered interface combinations. When considering only structures that exhibit the same interface combination, the distributions are markedly sharper than in figure 1 B, where the distributions were estimated from all structures. Thus, it seems likely that the structural complexity indicated by the distributions in the bound state is a result of a superposition of narrower distributions that belong to certain interface combinations (in addition to noise). A similar conformational behavior can also be observed in the simulation of the bivalent SIM2-SIM3-diSUMO3 complex in the overall parallel configuration (figure S13).

It is not surprising that structures with interface combination 2/1 are conformationally least restrained, as SIM2 cluster 2 comprises of structures with a single side chain in the SIM binding groove. Intuitively, this grants the complex more freedom and is thus entropically favorable. This may also be the reason for the relatively high probability of this interface combination, despite the fact that the individual SIM2-SUMO3 interface does not seem favorable in comparison to the typical SIM-SUMO interface.

From this point of view it is quite remarkable that interface combination 1/3 has relatively high probability and at the same time, structures with this interface combination are strongly conformationally restrained. The reason for this is strong electrostatic interaction (see table 3). On the one hand this likely stabilizes the 1/3 interface combination, on the other hand it also strongly restricts the conformational freedom of diSUMO3. As figure 1 C shows, electrostatic interactions between negatively charged residues of the linker between the SIMs and positively charged residues of both SUMO3 monomers are established that fix the structure of the complex.

**Table 3:** Average numbers of contacts between Asp/Glu of diSIM and Lys/Arg of diSUMO for each cluster. The column “all” simply gives the average of contacts between all those residues, the column “SIM2” only considers Asp4, Glu5 and Asp8. The “SIM3” column only considers Asp20 and Asp25 and the column “linker” only considers Glu12 and Glu15.

| clusters | all | SIM2 | SIM3 | linker |
| --- | --- | --- | --- | --- |
| SIM2 3/SIM3 2 | 2.56 | 0.19 | 1.56 | 0.81 |
| SIM2 1/SIM3 3 | 5.47 | 1.50 | 0.23 | 3.74 |
| SIM2 2/SIM3 1 | 3.76 | 1.65 | 0.58 | 1.53 |
| SIM2 4/SIM3 1 | 3.60 | 1.22 | 0.74 | 1.64 |

To investigate the behavior of the SIM2-SIM3 peptide in the bivalent complex relative to the unbound state of the peptide, we conducted an additional REST2 simulation of the free SIM2-SIM3 peptide in a temperature range from 303.15 K to 390 K and with a simulation time of 1 *μ*s per replica.

In figure S8 A, we consider the distance distributions of several relevant pairs of amino acids of the SIM2-SIM3 peptide and the distribution of the distance of the SIM binding grooves of proximal and distal SUMO3 in the bound and the free state. All considered distributions are very broad, both in the free and in the bound state and clearly overlap.

In figure S8 B, we consider the distributions of distances of amino acid pairs, estimated from structures that exhibit interface combinations in which these pairs occupy the SIM binding grooves. For example, we consider the distribution of the Leu9-Val19 distance estimated from the structures with interface combination 3/2, where these two residues occupy the SIM binding grooves of proximal and distal SUMO3. For all considered interface combinations, except for combination 2/1, these distributions are markedly sharper than those that were estimated from all bound structures. Interestingly, we observe that the distance distribution of the amino acid pair with the largest spacing, Ile6 and Val19, which occupy the two SIM binding grooves in interface combination 1/3, has a sharp peak at the smallest distance among all considered pairs. From this we conclude that the large spacing between these residues is not the reason for the high probability of interface combination 1/3, where these residues occupy the SIM binding grooves.

## Discussion

Our simulations aim to describe the possible structure of bivalent diSIM-diSUMO complexes at the example of the SIM2-SIM3 peptide of RNF4 and diSUMO3. They question previously reported models of complexes formed by SUMO chains and disordered peptides with multiple SIMs, in which the SUMO chains adopt a defined conformation or are at least strongly conformationally restrained in comparison to the unbound state. In fact, only one such model is fully experimental in nature, namely the NMR structure of the complex formed by RNF4-SIM2-SIM3 and diSUMO3 (PDB: 2mp2).^13^

In contrast to this NMR structure, our simulations surprisingly draw a much more complex picture of the diSUMO3 RNF4-SIM2-SIM3 interaction. Most importantly, there appears to be an equilibrium between relevant diSUMO3 conformations and also between different binding modes of the SIM regions. The known flexibility of free SUMO chains, the disordered nature of the RNF4-SIM region, the unspecific and non-essential contribution of the linker between the SIMs^11–14,27^ and the ability of SIMs to individually bind SUMO in two different orientations,^21,31^ however, all would reasonably support a structural model that would encompass a larger number of possible conformations as suggested by our analysis. Such a behavior with multiple conformations in the bound state has, for example, also been observed in bivalent complexes involving diUbiquitin.^26^ Importantly, in these complexes, di-Ubiquitin retained some conformational freedom despite the fact that the considered binding partners were far more rigid than the SIM region of RNF4.

We observed multiple relevant interfaces between the individual SIM regions and SUMO3 domains. Among these are the expected classical parallel and antiparallel SIM2- and SIM3-SUMO3 interfaces, but also other typical SIM-SUMO interfaces established by other residues of the SIM region. Interestingly, we furthermore observed interfaces that deviate from the typical SIM-SUMO interface, but were clearly relevant according to our analysis. These are intuitively unfavorable, but appear to be stabilized by electrostatic interaction of the linker region between the SIMs (SIM2 interface 1) or entropic effects (SIM2 interface 2).

In figure 4, we sketch an equilibrium of states of the RNF4-SIM-tetraSUMO complex with varying numbers of typical SIM-SUMO interfaces. Previously, it was assumed that this complex exhibits three or four SIM-SUMO interfaces in equilibrium, established by the classical SIMs of RNF4.^11–13^ Extending our observations on the bivalent complex to the tetravalent complex, it seems likely that in equilibrium also conformational states with a lower number of typical SIM-SUMO interfaces have relatively high probability. Furthermore, interfaces could also be established by other regions of the peptide than the four classical SIMs of RNF4. Such alternative states likely benefit from a higher conformational entropy of the SUMO chain.

**Figure 4:**
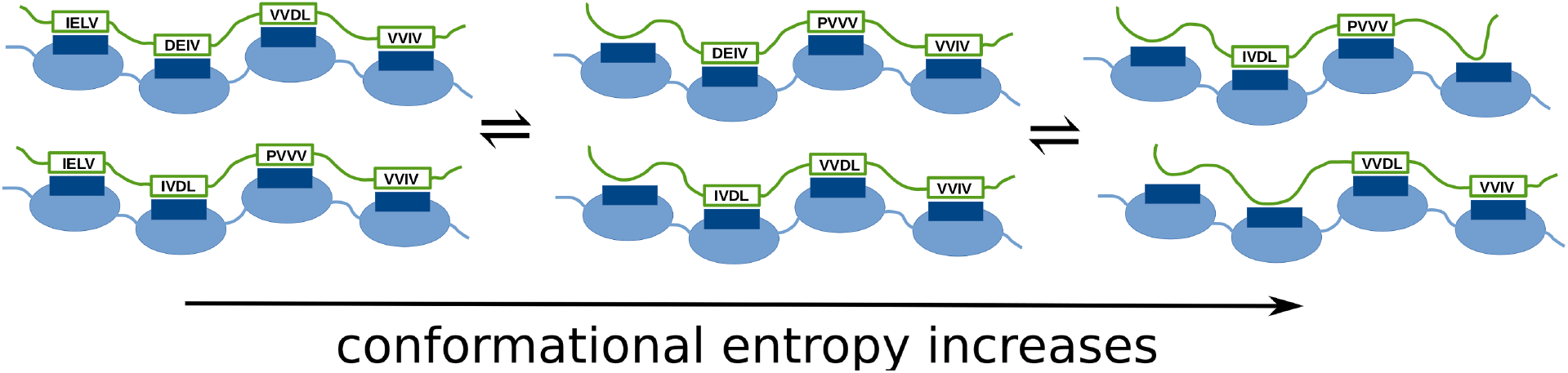
Equilibrium of states with two, three and four typical SIM-SUMO interfaces for the RNF4-SIM-tetraSUMO complex. SUMO monomers are represented by light blue elipses and their SIM binding groove by dark blue rectangles. Regions of the peptide that engage in a typical SIM-SUMO interface are depicted as green boxes with the residues that interact in them. Besides the typical SIM-SUMO interfaces, other interfaces may be established by regions of the peptide, similar to that of SIM2 cluster 2 in our simulations.

Keusekotten et al. measured the affinities of various mutants of the RNF4-SIM region and tetraSUMO2. ^12^ We summarize their results in table 4. A model in which only states with the typical SIM-SUMO interfaces of at least SIM2, SIM3 and SIM4 have high probability is clearly inconsistent with this data, as here the mutation of one of these SIMs would lead to a significant decrease in the affinity. A model based on our simulation data, in which multiple different complex configurations with varying interfaces are populated in equilibrium is, however, consistent with the data. Here, upon mutation of one of these SIMs, only a small subset of the states would be affected, thus there would only be a minor effect on the overall affinity, which is also what has been measured in the experiments.

**Table 4:**
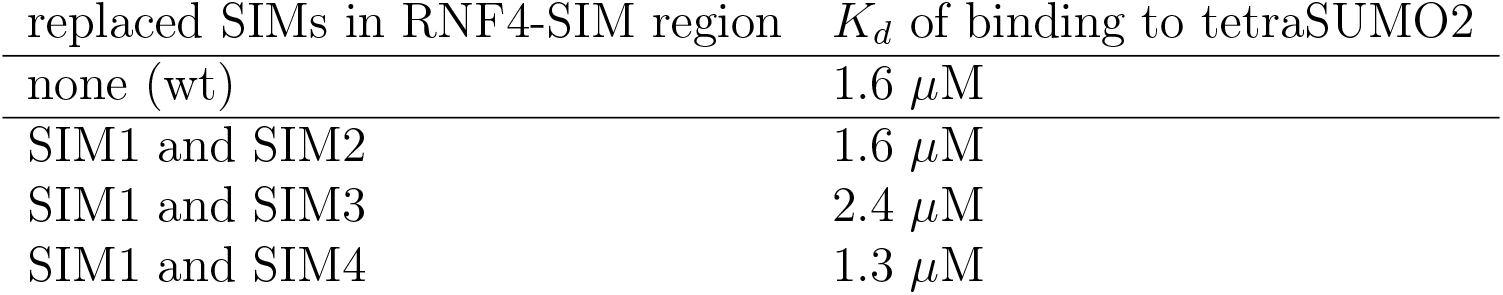
Dissociation constants of tetraSUMO2 and the RNF4-SIM region (residues 1-125) and mutants of it, where two SIMs have been replaced by residues AAAA, reported by Keusekotten et al. ^12^ Note that in case of SIM2 and SIM3 residues VDLT were mutated.

Our observation that residues outside the classical SIM2 and SIM3 of RNF4 can establish the interaction with the SIM binding groove of SUMO3 is also consistent with findings of a recent proteomics study. Here, a large number of proteins was identified as SUMO2 interaction partners that contain sequences with only remote similarity to the classical SIM consensus sequence.^25^

Typical SIMs of the type [V/I/L][V/I/L]DLT like SIM2 and SIM3 of RNF4 and in particular the SIM of PIAS2 (VIDLT) are sometimes considered optimized for the interaction with SUMO.^12,32^ Related to this is the concept of a dominant SIM that is much more important for the recognition of SUMO chains than other SIMs of a multi-SIM region. This concept was introduced by Sun et al. on the basis of mutation experiments of the SIM2 region of Arkadia/RNF111 (like SIM3 of RNF4, SIM2 of Arkadia has the sequence VVDLT).^32^ In this context, Sun et al. also found that residues Asp3, Leu4, and Thr5 of the SIM were strictly required for high affinity interaction with SUMO. This indicates that in the complex formed by Arkadia/RNF111 and SUMO chains, a single high probability SIM2-SUMO interface can be found, and that consequently, the binding modes of the two known human STUbLs RNF4 and Arkadia/RNF111 and SUMO chains differ.

Beside the interaction of regions with multiple SIMs and SUMO chains, also the interaction of such multiSIM regions and individual SUMO proteins that are conjugated to the same substrate is relevant. In this context, a preference of disordered SIM regions for SUMO chains was observed.^24^ It is expected that relative positions and orientations of multiple individually conjugated SUMOs on a substrate are rather fixed. Consequently, binding these SUMO proteins likely limits the conformational freedom of the disordered SIM region and is thus entropically unfavorable. Our simulations indicate that this is not the case for the interaction with SUMO chains. This likely contributes to the observed preference.

In our analysis and discussion, we often focused on the overall antiparallel configuration of the diSIM-diSUMO3 complex, as our simulations indicate that this might be more relevant. It is clear from the experiments of Xu et al., however, that the alternative, overall parallel arrangement of the complex is not irrelevant. Importantly, the central conclusions of the present study, namely the great interface variability and the conformational freedom of the SUMO3 dimer in complex with the SIM2-SIM3 peptide, are also valid for the overall parallel arrangement of the complex, as further detailed in figures S9-S13.

Our large scale simulations offer new insights into the nature of bivalent SIM-SUMO interaction and allowed us a reevaluation and new interpretation of previous experimental results. Thus our study also highlights the possible role of simulations as a complementary tool in the future investigation of the multivalent interaction with dynamic multidomain systems like SUMO and Ubiquitin chains.

## Methods

### Molecular dynamics simulations

We set up our systems using structures from PDB file 2mp2^13^ and processing them with the CHARMM-GUI.^33^ We used the CHARMM 36m force field^34^ and TIP3P water.^35^ We added sodium chloride in a concentration of 0.15 mM. We checked the protonation states of amino acids using propka integrated in the playmolecule web server. ^36^

We conducted all simulations using algorithms and parameters recommended for the use of the CHARMM force field and GROMACS. ^33^ In particular, we used Nosé-Hoover temperature coupling^37,38^ and Parrinello-Rahman pressure coupling^39^ for simulations at 303.15 K and atmospheric pressure. See reference 33 for more details.

We used Gromacs 2016.5^40^ patched with Plumed 2.4^41^ to conduct the REST2^28^ simulations, i.e. Hamiltonian replica exchange molecular dynamics.

REST2 is a variant of parallel tempering simulations. ^28^ The configuration space of systems like the presently considered SIM2-SIM3-diSUMO3 complex consists of multiple relevant states (i.e. structures) that are separated by energy barriers which are rarely crossed in conventional MD simulations. In parallel tempering, one exploits the fact that these energy barriers are crossed much faster at higher temperatures. One sets up multiple simulations in parallel that are conducted at increasing temperatures, the lowest typically being the one of interest. At certain time steps, one tries Monte Carlo exchanges of structures between adjacent temperatures. This way, at high temperatures the system explores the configuration space efficiently and the important low energy states trickle down to the temperature of interest. In REST2, instead of working with increasing temperature directly, one scales the interaction energy, thus working with effective temperatures. This has been shown to be more efficient than conventional parallel tempering.

For the plain MD simulations that were used to estimate dynamical properties, we conducted the simulations as just described, but used Gromacs version 2019 on GPUs.

For the REST2 simulations of the SIM2-SIM3-diSUMO3 complex, we performed the replica simulations at effective temperatures 303.15, 306.01, 309.02, 313.3, 317.81, 322.34, 327.21, 333.48, 339.86, 346.35, 352.95, 359.66, 366.48, 373.42, 380.47, 387.65, 394.96, 402.37, 409.93, 417.6, 425.41, 433.31, 441.39, 449.6, 457.94, 466.43, 475.06, 483.83, 492.76, 500. We do not assign a unit to these effective temperatures as they actually are scaling factors for the interaction of the protein and peptide atoms. The rate of successful exchanges between replicas was between 27 % and 58 %.

For the REST2 simulations of the free diSUMO3, we performed the replica simulations at effective temperatures 303.15, 306.01, 309.02, 313.3, 317.81, 322.34, 327.21, 333.48, 339.86, 346.35, 352.95, 359.66, 366.48, 373.42, 380.47, 387.65, 394.96. The rate of successful exchanges between replicas was between 32 % and 59 %.

Finally, for the REST2 simulations of the free SIM2-SIM3 peptide, we performed the replica simulations at effective temperatures 303.15, 315.1, 329.2, 346.99, 368.4, 390.91. The rate of successful exchanges between replicas was between 11 % and 27 %.

For the complex, a simulation time of 3 *μ*s per replica and for the free diSUMO3 and SIM2-SIM3 peptide 1 *μ*s per replica were used. This gives a combined simulation time of 203 *μ*s.

### Analysis

The cluster analysis was performed using the HDBSCAN algorithm. ^42,43^ For example, to identify relevant clusters of the SIM2 region, we first aligned the full structure of the complex with respect to the SUMO3 monomer bound to the SIM2 region. Subsequently, we subjected the positions of the nitrogen, C*α*, C*β* and carbonyl carbon atoms of several residues to the cluster algorithm. We identified SIM2 clusters 1 and 2 using residues Asp4 to Val7 of the SIM2-SIM3 peptide, SIM2 clusters 3 and 4 using residues Ile6 to Leu9 of the SIM2-SIM3 peptide, SIM3 clusters 1 and 2 using residues Pro16 to Val19 and SIM3 clusters 3 and 4 using residues Val18 to Leu21. We remark that no to structures were assigned two the same SIM2 or SIM3 cluster.

When considering the root mean squared deviation of atomic positions (RMSD) of structures within a cluster, we obviously did not perform an additional optimal alignment before computing the RMSD as is otherwise common. We calculated the RMSD for the same atoms that were used in the respective cluster analysis.

Similarly, we did not optimally align the structures of C-SUMO when investigating the diSUMO3 conformation (figure 3).

We frequently considered the most representative structure 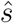 within some set of structures *S*:

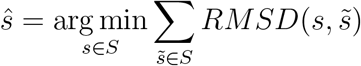

For the visualization of protein structures we used VMD. ^44^

