## Supporting Information for "Conformational and interface variability of bivalent RNF4-SIM diSUMO3 interaction"

June 15, 2021

The analysis in the main text focuses on the simulation started in the best representative conformer of PDB file 2mp2, in which the complex adopts an overall antiparallel orientation with interaction between the SIM2 region and the distal SUMO3 and the SIM3 region and the proximal SUMO3. Supporting figures and tables concerning this simulation are presented in the first two sections. Figures and tables concerning the simulation of the complex in the overall parallel orientation follow in the third and fourth section.

### Supporting figures for the overall antiparallel orientation

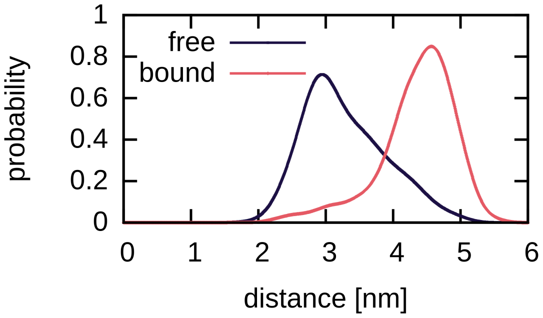


Figure 1: Distribution of the Arg60-Arg60 distance in the bound and the free state.

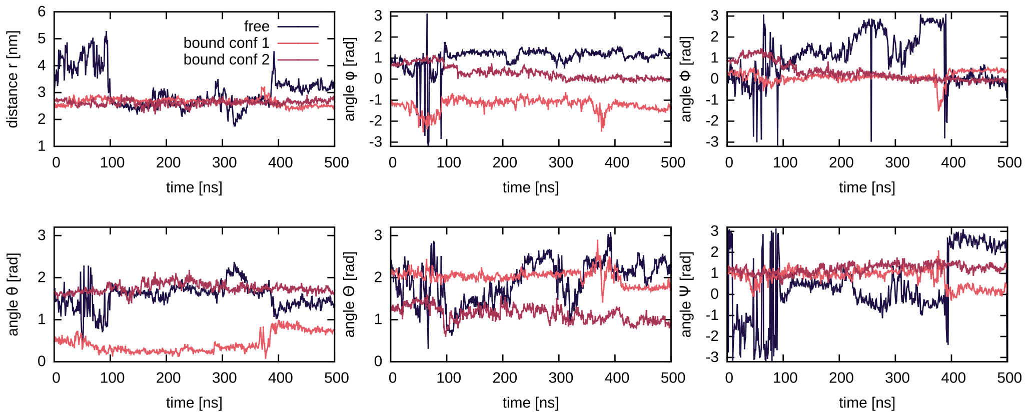


Figure 2: Time series of the collective variables defined in figure 1 B of the main text in the additional ordinary MD simulations of the bound and free systems. For the bound system two simulations were started from different initial structures (bound conf 1 and bound conf 2). For reasons of visualization, we shifted some of the time series by a constant value.

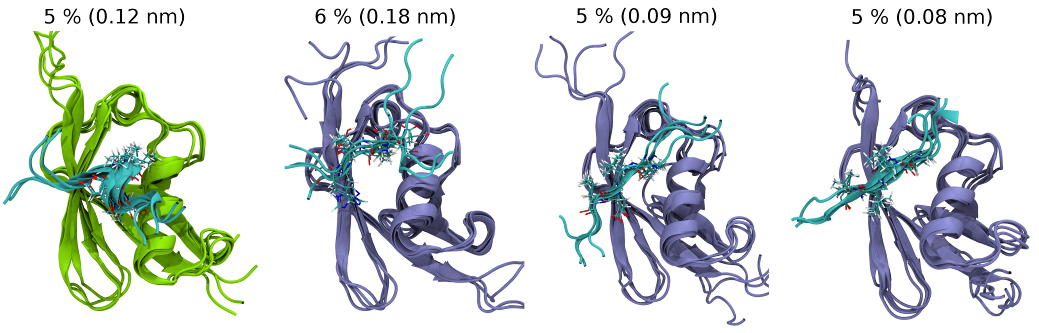


Figure 3: Representative structures for one additional cluster of the SIM2 region (left) and three additional clusters of the SIM3 regions. The probabilities of the clusters are given as percentages and the RMSD of the median structure and the most representative structure is given in parenthesis. In the cluster of the SIM2 region, the SIM2 residues form a helix. In the three clusters of the SIM3 region Leu21 occupies the SIM binding groove and the SIM region adopts an antiparallel orientation, Val18 and Asp20 occupy the SIM binding groove and the SIM region adopts an antiparallel orientation and Pro16 and Val18 occupy the SIM binding groove and again, the SIM region adopts an antiparallel orientation (second to forth structure).

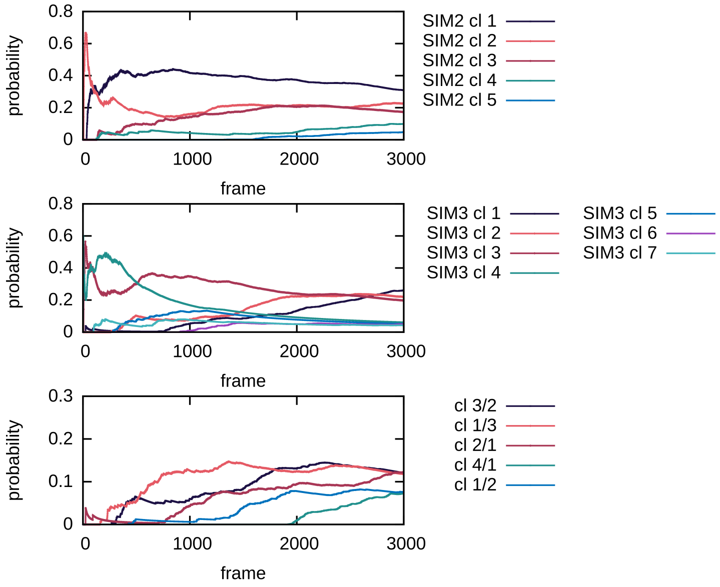


Figure 4: Simulation time dependence of the estimates for the cluster probabilities of the SIM2 region, the SIM3 region, and high probability combinations. We refer to the simulation time or equivalently the analyzed snapshots from the simulation as frame because the dynamical information is lost with the use of replica exchange.

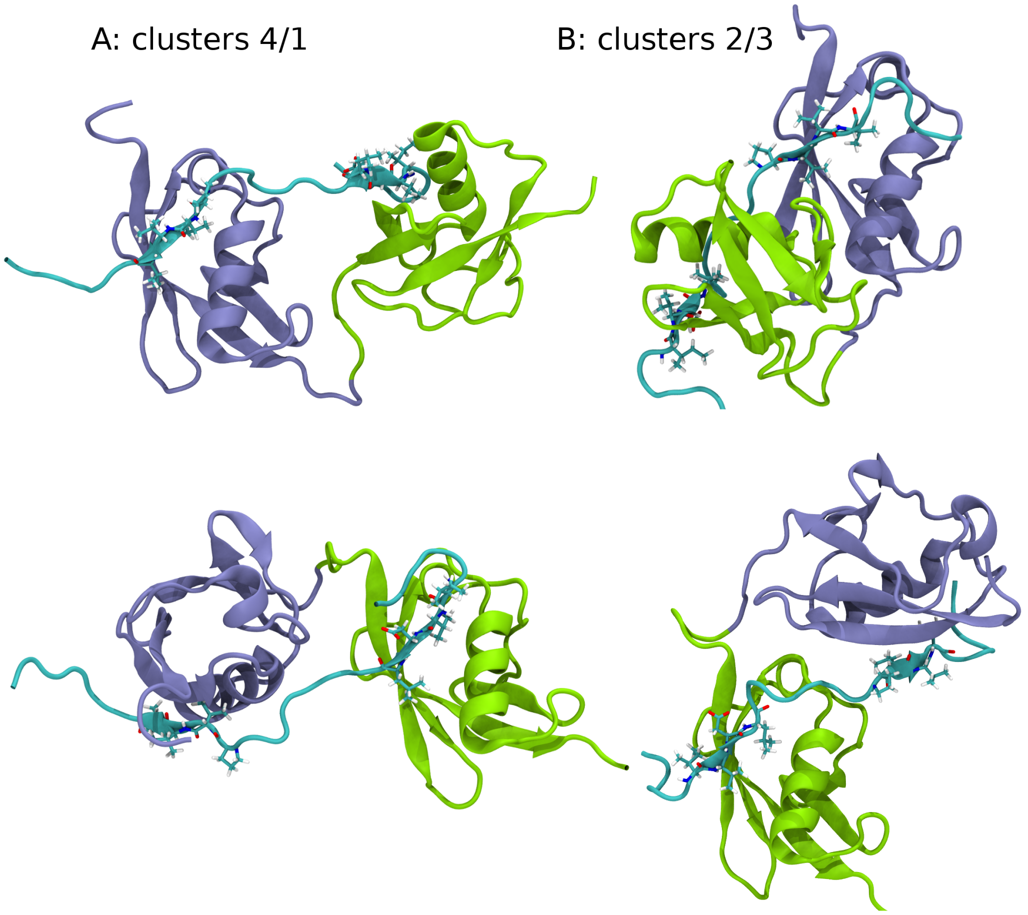


Figure 5: A: A structure of the complex with interface combination 4/1, i.e. the SIM2 interface falls in cluster 4 and the SIM3 interface falls in cluster 1. Here, both SIMs bind in a typical antiparallel SIM-SUMO interface. Top and bottom figure show the same structure from different perspectives. B: A structure of the complex with interface combination 3/2. Here, both SIMs bind in a typical parallel SIM-SUMO interface. Again, top and bottom figure show the same structure from different perspectives. Clearly, the relative position of the SUMO3 domains is different for the two structures. Distal (N-terminal) SUMO is depicted in violet, proximal (C-terminal) SUMO is depicted in green and the SIM2-SIM3-peptide is depicted in cyan.

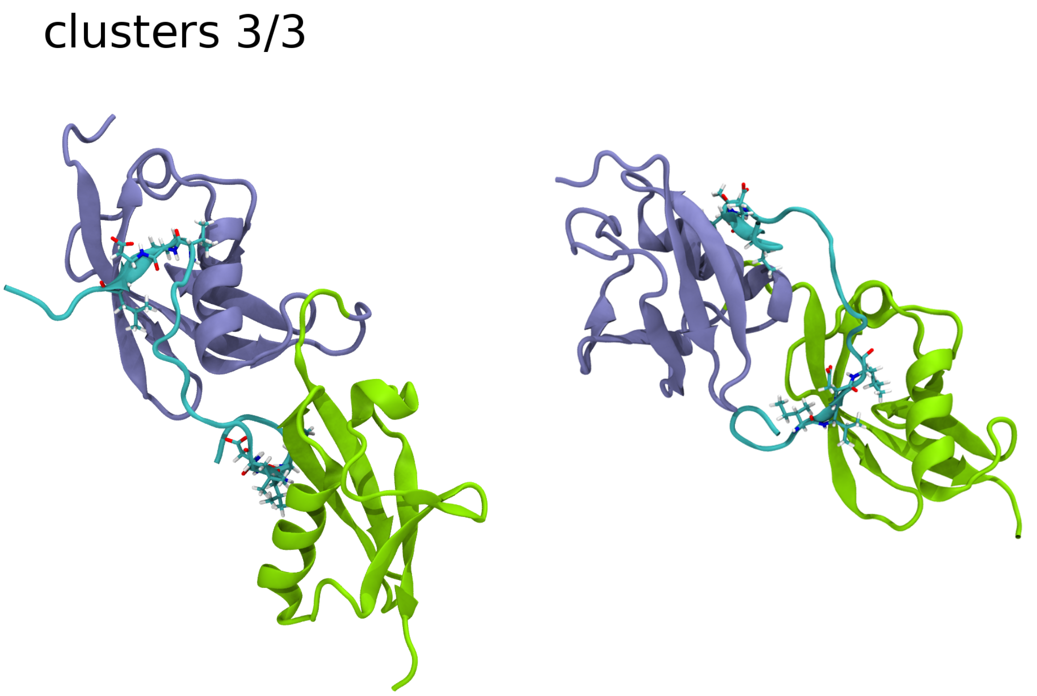


Figure 6: The figure shows a structure of the complex with interface combination 3/3, i.e. the SIM2 interface falls in cluster 3 and the SIM3 interface falls in cluster 3. Here, SIM2 binds in the typical parallel and SIM3 binds in the typical antiparallel SIM-SUMO interface. This interface combination has very low probability despite the presence of the typical SIM-SUMO interfaces. While the SIM3 region seem unfavorably bent, there do not seem to be obvious steric reasons for the proximal SUMO3 to not move up right relative to the distal SUMO3 an thus allow the SIM3 region to interact in a more favorable manner. We did not observe this in our simulations, however. Distal (N-terminal) SUMO is depicted in violet, proximal (C-terminal) SUMO is depicted in green and the SIM2-SIM3-peptide is depicted in cyan.

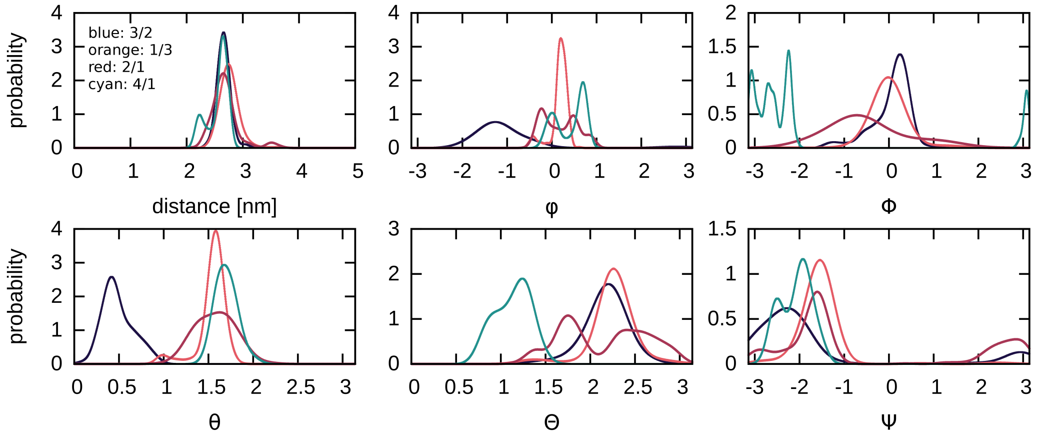

Figure 7: Distributions of collective variables that capture the diSUMO3 conformation, defined in figure 1 of the main text, for four interface combinations.

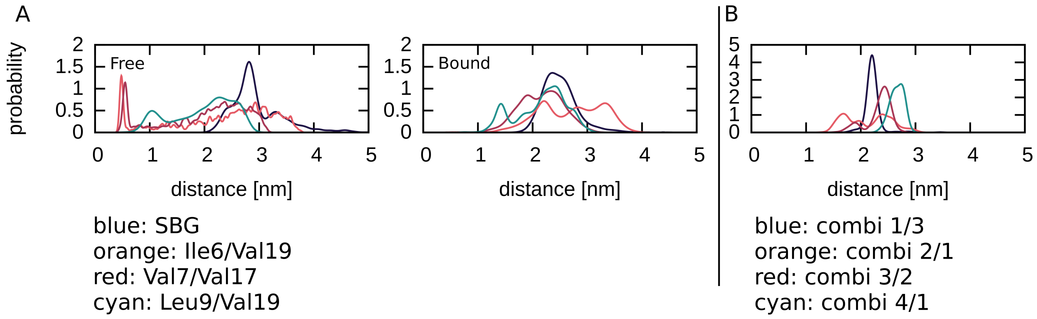


Figure 8: A: Distributions of the distances of several pairs of amino acids of the SIM2-SIM3 peptide and the SIM binding grooves of proximal and distal SUMO3 in the free and the bound state. For the SIM binding groove, we consider the center of residues Lys32, Ile33, Lys41 and Leu42 of SUMO3, this corresponds to the spot close to the first helix turn. B: Distributions of the of the same amino acid pairs, estimated only from structures with interface combination 1/3 (residues Ile6 and Val19), 2/1 (residues Val7 and Val17), 3/2 (residues Leu9 and Val19) and 4/1 (residues Val7 and Val17).

### Supporting tables for the overall antiparallel orientation

Table 1: Probabilities of combinations of low probability SIM3 interfaces and SIM2 interfaces. The product of the probabilities of the individual clusters is given in parenthesis.

| | SIM3 4 | SIM3 5 | SIM3 6 | SIM3 7 |
|---|---|---|---|---|
| SIM2 noise | 0.01 (0.01) | 0.00 (0.01) | 0.01 (0.01) | 0.02 (0.01) |
| SIM2 1 | 0.03 (0.02) | 0.05 (0.02) | 0.00 (0.02) | 0.00 (0.01) |
| SIM2 2 | 0.02 (0.01) | 0.00 (0.01) | 0.04 (0.01) | 0.00 (0.01) |
| SIM2 3 | 0.00 (0.01) | 0.00 (0.01) | 0.00 (0.01) | 0.02 (0.01) |
| SIM2 4 | 0.00 (0.01) | 0.00 (0.01) | 0.00 (0.01) | 0.00 (0.00) |
| SIM2 5 | 0.00 (0.00) | 0.00 (0.00) | 0.00 (0.00) | 0.00 (0.00) |

Table 2: Probabilities of combinations of high probability SIM3 interfaces and SIM2 interface 5. The product of the probabilities of the individual clusters is given in parenthesis.

| | SIM3 noise | SIM3 1 | SIM3 2 | SIM3 3 |
|---|---|---|---|---|
| SIM2 5 | 0.00 (0.01) | 0.05 (0.01) | 0.00 (0.01) | 0.00 (0.01) |

### Supporting figures for the overall parallel orientation

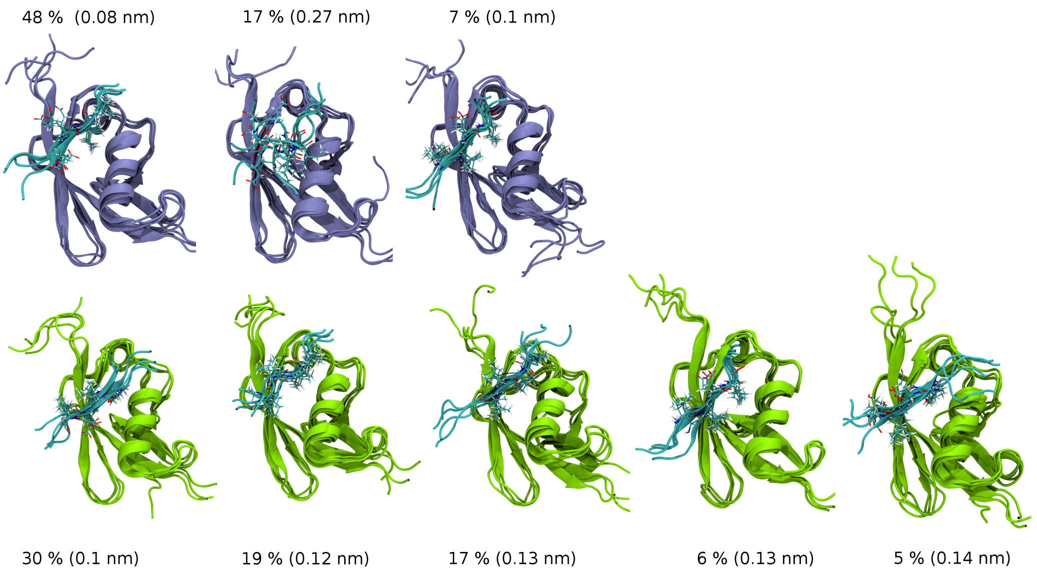


Figure 9: Representative structures for SIM2 clusters (top) and SIM3 clusters (bottom) in the overall parallel configuration. **SIM2 clusters from left to right:** Asp4 and Ile6 occupy the SIM binding groove and the SIM region forms a parallel $\beta$-sheet (similar to SIM2 cluster 1 of the overall antiparallel configuration), only Val7 occupies the SIM binding groove (similar to SIM2 cluster 2 of the overall antiparallel configuration) and Val7 and Leu9 occupy the SIM binding groove in the typical parallel SIM2-SUMO interface. **SIM3 clusters from left to right:** Val18 and Asp20 occupy the SIM binding groove and the SIM3 region forms an antiparallel $\beta$-sheet (similar to SIM3 cluster 5 of the overall antiparallel configuration). In the next cluster Pro16 and Val18 occupy the SIM binding groove in parallel orientation, the SIM3 region does not form a $\beta$-sheet however. In the next cluster, Val17 and Val19 occupy the SIM binding groove and the SIM3 region forms an antiparallel $\beta$-sheet (similar to SIM3 cluster 1 of the overall antiparallel configuration). The last two clusters of the SIM3 region comprise of the typical SIM3-SUMO interfaces in antiparallel and parallel orientation, respectively. See also table 3

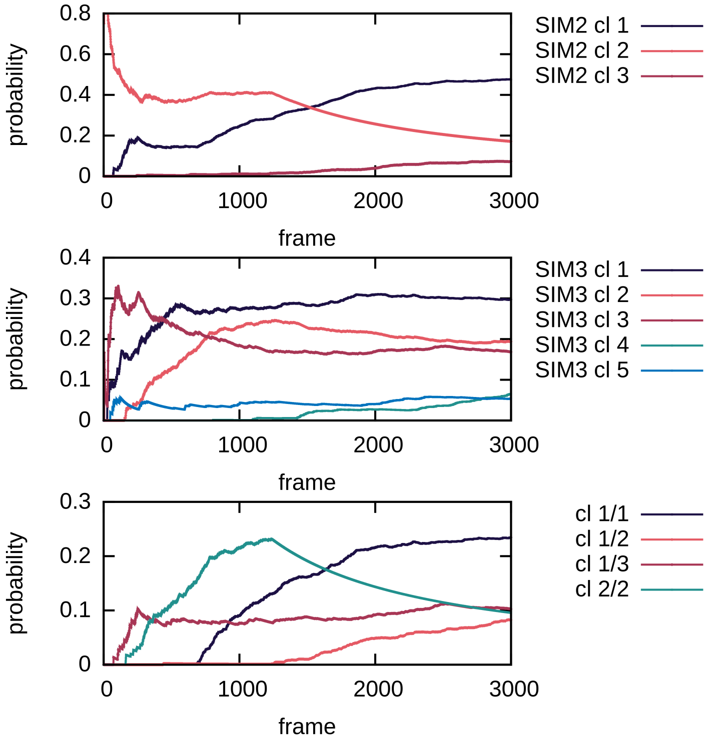


Figure 10: Simulation time dependence of the estimates for the cluster probabilities of the SIM2 region, the SIM3 region, and high probability combinations. We refer to the simulation time or equivalently the analyzed snapshots from the simulation as frame because the dynamical information is lost with the use of replica exchange.

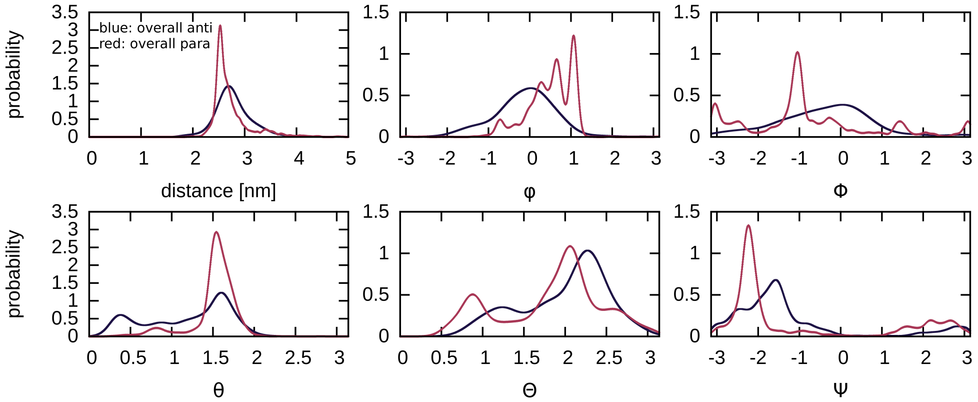


Figure 11: Comparison of the distributions of collective variables that capture the diSUMO3 conformation between overall antiparallel orientation (blue) and overall parallel orientation (red).

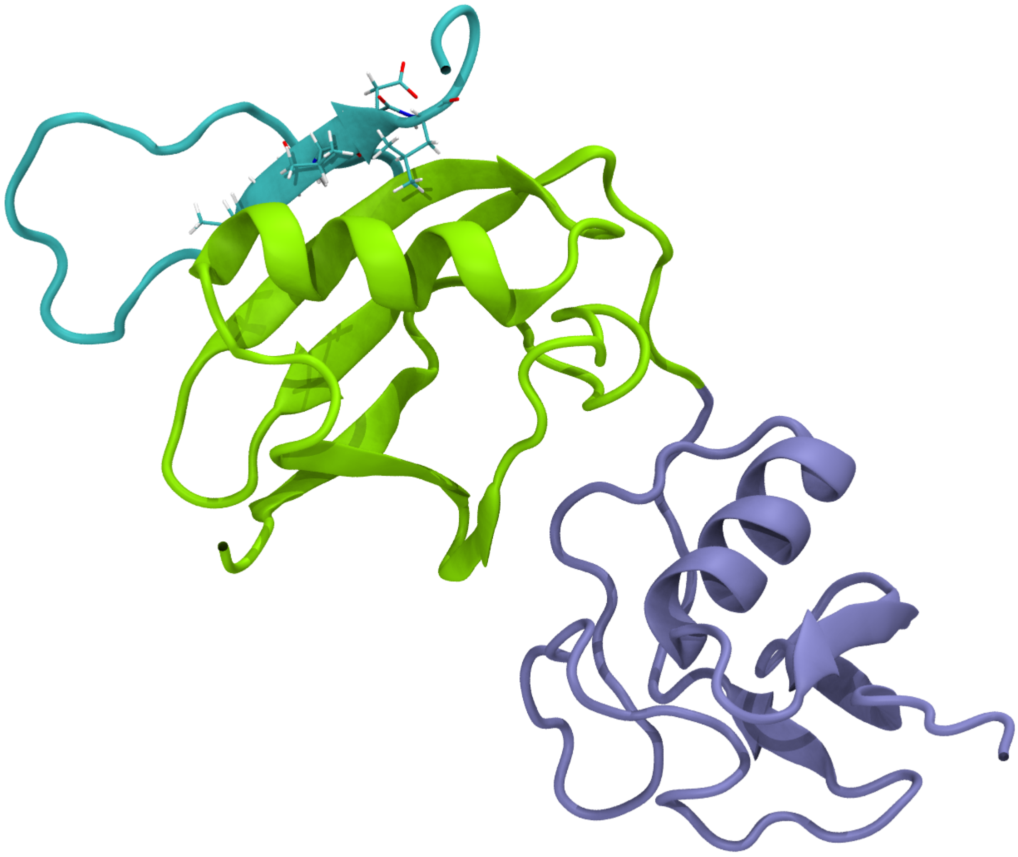

Figure 12: In this structure, only the SIM3 region interacts with a SUMO3 domain through a typical SIM3-SUMO interface.

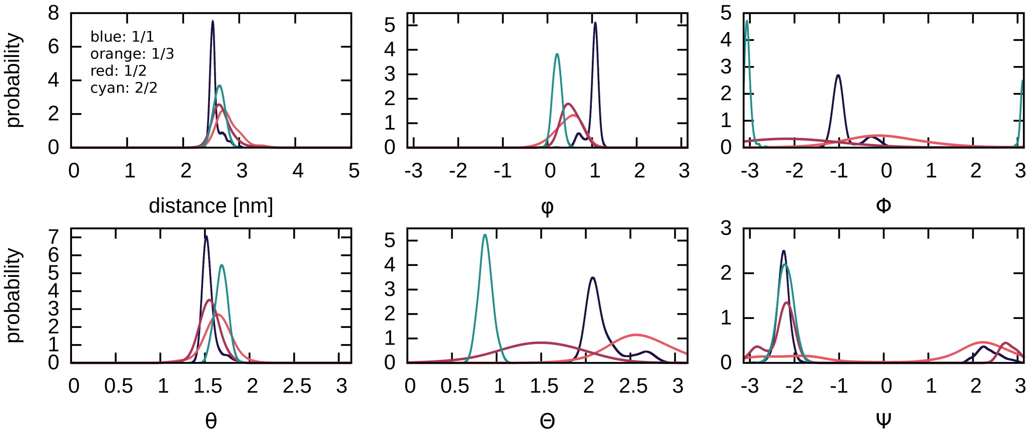


Figure 13: Distributions of collective variables that capture the diSUMO3 conformation, defined in figure 1 of the main text, for four interface combinations.

### Supporting tables for the overall parallel orientation

Table 3: Results of the cluster analysis: The column "SBG residues" and "orientation" give the residues that occupy the SIM binding groove and the orientation of the SIM region with respect to the SUMO3 monomer it binds to. $p_{tot}$, $p_{1st}$, $p_{2nd}$ and $p_{3rd}$ refer to the probabilities of the clusters estimated from the full simulation and from the first, second and third 1000 ns.

| cluster | SBG residues | orientation | $p_{tot}$ | $p_{1st}$ | $p_{2nd}$ | $p_{3rd}$ |
|---|---|---|---|---|---|---|
| SIM2 1 | Asp4, Ile6 | para | 0.25 | 0.62 | 0.57 | 0.48 |
| SIM2 2 | Val7 | para | 0.41 | 0.11 | 0 | 0.17 |
| SIM2 3 | Val7 Leu9 | para | 0.01 | 0.07 | 0.14 | 0.07 |
| SIM3 1 | Val18 Asp20 | anti | 0.27 | 0.35 | 0.27 | 0.30 |
| SIM3 2 | (Pro16) Val18 | para | 0.23 | 0.20 | 0.15 | 0.19 |
| SIM3 3 | Val17 Val19 | anti | 0.19 | 0.15 | 0.17 | 0.17 |
| SIM3 4 | Val19 Leu21 | anti | 0.00 | 0.10 | 0.23 | 0.06 |
| SIM3 5 | Val19 Leu21 | para | 0.04 | 0.04 | 0.08 | 0.05 |

Table 4: Probabilities of interface combinations. The product of the probabilities of the individual clusters is given in parenthesis. We underlined some interface combinations with relatively high probability. SIM2 n and SIM3 n refer to noise, i.e. structures that were not assigned to a cluster.

| | SIM3 n | SIM3 1 | SIM3 2 | SIM3 3 | SIM3 4 | SIM3 5 |
|---|---|---|---|---|---|---|
| SIM2 n | 0.09 (0.06) | 0.06 (0.08) | 0.02 (0.05) | 0.03 (0.05) | 0.06 (0.02) | 0.01 (0.01) |
| SIM2 1 | 0.05 (0.11) | 0.23 (0.14) | 0.08 (0.09) | 0.10 (0.08) | 0.00 (0.03) | 0.01 (0.02) |
| SIM2 2 | 0.07 (0.04) | 0.00 (0.05) | 0.10 (0.03) | 0.00 (0.03) | 0.00 (0.01) | 0.00 (0.01) |
| SIM2 3 | 0.02 (0.02) | 0.00 (0.02) | 0.00 (0.01) | 0.03 (0.01) | 0.00 (0.00) | 0.04 (0.00) |
